## Supplemental File for "Pyruvate and Related Energetic Metabolites Modulate Resilience Against High Genetic Risk for Glaucoma"

### Supplementary File

**Supplementary Methods:**

A total of 168 metabolites were quantified using nuclear magnetic resonance (NMR) spectroscopy. The panel includes Total Cholesterol, Total Cholesterol Minus HDL-C, Remnant Cholesterol (Non-HDL, Non-LDL Cholesterol), VLDL Cholesterol, Clinical LDL Cholesterol, LDL Cholesterol, HDL Cholesterol, Total Triglycerides, Triglycerides in VLDL, Triglycerides in LDL, Triglycerides in HDL, Total Phospholipids in Lipoprotein Particles, Phospholipids in VLDL, Phospholipids in LDL, Phospholipids in HDL, Total Esterified Cholesterol, Cholesteryl Esters in VLDL, Cholesteryl Esters in LDL, Cholesteryl Esters in HDL, Total Free Cholesterol, Free Cholesterol in VLDL, Free Cholesterol in LDL, Free Cholesterol in HDL, Total Lipids in Lipoprotein Particles, Total Lipids in VLDL, Total Lipids in LDL, Total Lipids in HDL, Total Concentration of Lipoprotein Particles, Concentration of VLDL Particles, Concentration of LDL Particles, Concentration of HDL Particles, Average Diameter for VLDL Particles, Average Diameter for LDL Particles, Average Diameter for HDL Particles, Phosphoglycerides, Total Cholines, Phosphatidylcholines, Sphingomyelins, Apolipoprotein B, Apolipoprotein A1, Total Fatty Acids, Degree of Unsaturation, Omega-3 Fatty Acids, Omega-6 Fatty Acids, Polyunsaturated Fatty Acids, Monounsaturated Fatty Acids, Saturated Fatty Acids, Linoleic Acid, Docosahexaenoic Acid, Alanine, Glutamine, Glycine, Histidine, Total Concentration of Branched-Chain Amino Acids (Leucine + Isoleucine + Valine), Isoleucine, Leucine, Valine, Phenylalanine, Tyrosine, Glucose, Lactate, Pyruvate, Citrate, 3-Hydroxybutyrate, Acetate, Acetoacetate, Acetone, Creatinine, Albumin, Glycoprotein Acetyls, Concentration of Chylomicrons and Extremely Large VLDL Particles, Total Lipids in Chylomicrons and Extremely Large VLDL, Phospholipids in Chylomicrons and Extremely Large VLDL, Cholesterol in Chylomicrons and Extremely Large VLDL, Cholesteryl Esters in Chylomicrons and Extremely Large VLDL, Free Cholesterol in Chylomicrons and Extremely Large VLDL, Triglycerides in Chylomicrons and Extremely Large VLDL, Concentration of Very Large VLDL Particles, Total Lipids in Very Large VLDL, Phospholipids in Very Large VLDL, Cholesterol in Very Large VLDL, Cholesteryl Esters in Very Large VLDL, Free Cholesterol in Very Large VLDL, Triglycerides in Very Large VLDL, Concentration of Large VLDL Particles, Total Lipids in Large VLDL, Phospholipids in Large VLDL, Cholesterol in Large VLDL, Cholesteryl Esters in Large VLDL, Free Cholesterol in Large VLDL, Triglycerides in Large VLDL, Concentration of Medium VLDL Particles, Total Lipids in Medium VLDL, Phospholipids in Medium VLDL, Cholesterol in Medium VLDL, Cholesteryl Esters in Medium VLDL, Free Cholesterol in Medium VLDL, Triglycerides in Medium VLDL, Concentration of Small VLDL Particles, Total Lipids in Small VLDL, Phospholipids in Small VLDL, Cholesterol in Small VLDL, Cholesteryl Esters in Small VLDL, Free Cholesterol in Small VLDL, Triglycerides in Small VLDL, Concentration of Very Small VLDL Particles, Total Lipids in Very Small VLDL, Phospholipids in Very Small VLDL, Cholesterol in Very Small VLDL, Cholesteryl Esters in Very Small VLDL, Free Cholesterol in Very Small VLDL, Triglycerides in Very Small VLDL, Concentration of IDL Particles, Total Lipids in IDL, Phospholipids in IDL, Cholesterol in IDL, Cholesteryl Esters in IDL, Free Cholesterol in IDL, Triglycerides in IDL, Concentration of Large LDL Particles, Total Lipids in Large LDL, Phospholipids in Large LDL, Cholesterol in Large LDL, Cholesteryl Esters in Large LDL, Free Cholesterol in Large LDL, Triglycerides in Large LDL, Concentration of Medium LDL Particles, Total Lipids in Medium LDL, Phospholipids in

Medium LDL, Cholesterol in Medium LDL, Cholesteryl Esters in Medium LDL, Free Cholesterol in Medium LDL, Triglycerides in Medium LDL, Concentration of Small LDL Particles, Total Lipids in Small LDL, Phospholipids in Small LDL, Cholesterol in Small LDL, Cholesteryl Esters in Small LDL, Free Cholesterol in Small LDL, Triglycerides in Small LDL, Concentration of Very Large HDL Particles, Total Lipids in Very Large HDL, Phospholipids in Very Large HDL, Cholesterol in Very Large HDL, Cholesteryl Esters in Very Large HDL, Free Cholesterol in Very Large HDL, Triglycerides in Very Large HDL, Concentration of Large HDL Particles, Total Lipids in Large HDL, Phospholipids in Large HDL, Cholesterol in Large HDL, Cholesteryl Esters in Large HDL, Free Cholesterol in Large HDL, Triglycerides in Large HDL, Concentration of Medium HDL Particles, Total Lipids in Medium HDL, Phospholipids in Medium HDL, Cholesterol in Medium HDL, Cholesteryl Esters in Medium HDL, Free Cholesterol in Medium HDL, Triglycerides in Medium HDL, Concentration of Small HDL Particles, Total Lipids in Small HDL, Phospholipids in Small HDL, Cholesterol in Small HDL, Cholesteryl Esters in Small HDL, Free Cholesterol in Small HDL, and Triglycerides in Small HDL. A subset of 27 metabolites certified for broad diagnostic use by the European Union (EU) includes Total cholesterol, VLDL cholesterol, Clinical LDL cholesterol, HDL cholesterol, Total triglycerides, Apolipoprotein B, Apolipoprotein A1, Total fatty acids, Omega-3 fatty acids, Omega-6 fatty acids, Polyunsaturated fatty acids, Monounsaturated fatty acids, Saturated fatty acids, Docosahexaenoic acid, Alanine, Glycine, Histidine, Total concentration of branched-chain amino acids, Isoleucine, Leucine, Valine, Phenylalanine, Tyrosine, Glucose, Creatinine, Albumin, and Glycoprotein acetyls.

**Supplemental Table 1. Unadjusted levels of plasma metabolites lactate, pyruvate, and citrate in UK Biobank participants and among the top 10% of glaucoma polygenic risk score (PRS)**

| | Total population | | | | Top 10% PRS | | | |
|---|---|---|---|---|---|---|---|---|
| **Characteristic** | **Overall** | **No Glaucoma** | **Glaucoma** | **p-value** | **Overall** | **No Glaucoma** | **Glaucoma** | **P-value** |
| Sample size | 117,698 | 113,040 | 4,658 | | 11,770 | 10,077 | 1,693 | |
| Lactate, mmol/l (median [IQR]) | 3.94 [3.24, 4.75] | 3.95 [3.24, 4.76] | 3.82 [3.17, 4.60] | <0.001 | 3.94 [3.25, 4.75] | 3.96 [3.27, 4.78] | 3.79 [3.16, 4.58] | <0.001 |
| Pyruvate, mmol/l (median [IQR]) | 0.080 [0.063, 0.10] | 0.080 [0.06, 0.10] | 0.0780 [0.06, 0.10] | <0.001 | 0.080 [0.06, 0.10] | 0.080 [0.06, 0.10] | 0.077 [0.06, 0.09] | <0.001 |
| Citrate, mmol/l (median [IQR]) | 0.065 [0.057, 0.074] | 0.065 [0.06, 0.07] | 0.065 [0.06, 0.07] | 0.51 | 0.065 [0.06, 0.07] | 0.06 [0.06, 0.07] | 0.07 [0.06, 0.07] | 0.87 |

Abbreviations IQR, interquartile range

**Supplemental Table 2. Metabolite data beta (effect size) values by model and metabolite groupings.**

See excel supplemental. Model 1 includes metabolites only; Model 2 incorporates additional covariates including age, sex, genetic ancestry, season, time of day of specimen collection, and fasting time; Model 3 incorporates covariates in Model 2 and smoking status (never, past, and current smoker), alcohol intake (g/week), caffeine intake (mg/day), physical activity (metabolic equivalent of task [MET], hours/week), body mass index (kg/m$^2$), average systolic blood pressure (mm Hg), history of diabetes, HbA1c (mmol/mol), history of coronary artery disease, systemic beta-blocker use, oral steroid use, and spherical equivalent refractive error (diopters); Model 4 incorporates covariates in model 3 and a glaucoma polygenic risk score (PRS). Each color represents a different panel of metabolites (grey = no metabolites; light blue = 27 metabolites; and dark blue = 168 metabolites). The white text represents the AUC ± 95% confidence interval. Abbreviations: EU, European Union.

**Supplemental Table 3. Metabolic risk score beta (effect size) values.**
See excel supplemental.

**A.**

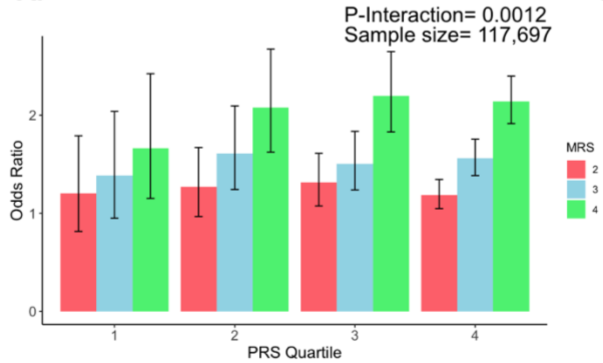


| | Quartiles | | | |
|---|---|---|---|---|
| | Q1 | Q2 | Q3 | Q4 |
| **PRS** | -0.0090, -0.0075 | -0.0075, -0.0073 | -0.0073, -0.0070 | -0.0070, -0.0055 |
| **MRS** | 0.0000, 0.0316 | 0.0316, 0.0384 | 0.0384, 0.0462 | 0.0462, 0.5185 |

**B.**

| | Glaucoma cases | Glaucoma Metabolite Reference | Odds Ratio (95% CI) | p-value |
|---|---|---|---|---|
| **PRS Q1** | | | | |
| MRS Q2 | 56 | 46 | 1.20 (0.81, 1.79) | 0.40 |
| MRS Q3 | 64 | 46 | 1.39 (0.95, 2.04) | 0.11 |
| MRS Q4 | 73 | 46 | 1.66 (1.15, 2.42) | 0.0086 |
| **PRS Q2** | | | | |
| MRS Q2 | 119 | 94 | 1.27 (0.97, 1.67) | 0.098 |
| MRS Q3 | 149 | 94 | 1.61 (1.24, 2.09) | 0.0004 |
| MRS Q4 | 192 | 94 | 2.08 (1.62, 2.67) | <0.0001 |
| **PRS Q3** | | | | |
| MRS Q2 | 220 | 171 | 1.31 (1.07, 1.61) | 0.0091 |
| MRS Q3 | 254 | 171 | 1.51 (1.24, 1.83) | <0.0001 |
| MRS Q4 | 365 | 171 | 2.20 (1.83, 2.65) | <0.0001 |
| **PRS Q4** | | | | |
| MRS Q2 | 582 | 495 | 1.19 (1.05, 1.34) | 0.0077 |
| MRS Q3 | 747 | 495 | 1.56 (1.39, 1.76) | <0.0001 |
| MRS Q4 | 1031 | 495 | 2.14 (1.91, 2.40) | <0.0001 |

**Supplemental Figure 1. Interaction of the holistic metabolite risk score (n=168 metabolites) and polygenic risk score (PRS) on glaucoma risk** (A) The bar chart shows the interaction of holistic probit-transformed metabolite risk score (MRS) with glaucoma genetic predisposition in each PRS quartile. In each glaucoma PRS quartile, the lowest metabolic sum quartile (Q1) is the metabolite reference group used to calculate the odds ratios. Each color represents the MRS quartiles (red = second quartile; blue = third quartile; and green = fourth quartile). Error bars show 95% confidence interval (CI). The table under the bar chart shows the ranges for the PRS and MRS quartiles. (B) Table showing odds ratios for glaucoma by polygenic risk score (PRS) and MRS within various quartiles. The number of glaucoma cases within each MRS quartile and the number of glaucoma cases in the first quartile of MRS (Q1), labeled as glaucoma metabolite reference, are used to calculate the odds ratios. This analysis is adjusted for time since the last meal/drink (hours), age, age-squared, sex, ethnicity (Asian, Black, White, and other), season, time of day of specimen collection (morning, afternoon, night), smoking status (never, past, and current smoker), alcohol intake, caffeine intake, physical activity (metabolic equivalent of task [MET] hours/week), body mass index (kg/m$^2$), average systolic blood pressure (mm Hg), history of diabetes (yes or no), HbA1c (mmol/mol), history of coronary artery disease, systemic beta-blocker use, oral steroid use, and spherical equivalent refractive error (diopters).

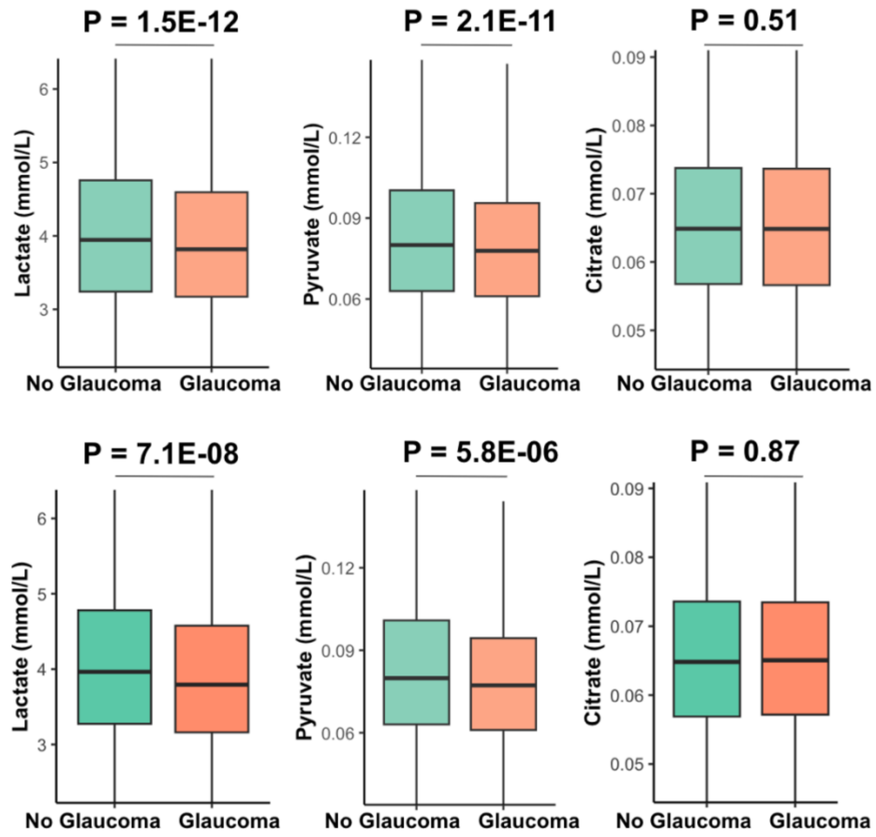


**Supplemental Figure 2. Unadjusted levels of plasma metabolites lactate, pyruvate, and citrate in all participants (top row) and among the top 10% of glaucoma polygenic risk score (bottom row).**
